# Odor sequence learning in honeybees: insights from olfactory classical conditioning paradigm

**DOI:** 10.1101/2025.07.04.663174

**Authors:** Maria Bortot, Giorgio Vallortigara

## Abstract

The ability to extract regularities from the sensory input is important for adaptation to a complex environment and to support behavioral flexibility. Honeybees have sophisticated cognitive and learning capacities, including categorization of stimuli on the basis of perceptual (e.g., symmetry) and abstract (e.g., sameness/difference) concepts. Here we investigated whether bees could extrapolate the temporal regularity of a sequence. By exploiting the olfactory conditioning of the proboscis extension response (PER), we investigated whether honeybees could learn and generalize a particular sequence structure composed of different odors presented in a specific temporal order. Different conditioning paradigms (i.e., absolute, differential, generalization) were employed. Bees used different strategies, such as an early tendency to encode the single odor properties, instead of learning the entire sequence pattern. The use of a generalization paradigm potentially uncovered a spontaneous tendency to transfer over similar structures one hour after the training irrespective of the single-element properties, though this finding is not yet conclusive. These results shed light on the strategies used by bees to solve an odor abstraction task, highlighting the crucial role of the type of conditioning to let them emerge.

## Introduction

From an evolutionary perspective, inferring a rule that allows the treatment of different stimuli as belonging to the same category could be extremely important. It might help an animal in anticipating a particular outcome, as well as to generalize its response, thus maximizing its fitness. This capacity has been thought to be one of the core mechanisms underlying human language acquisition, albeit not limited to our species [1].

The ability to extrapolate regularities and generalize them to new stimuli has been found in different vertebrate species, such as primates and birds [2–5]. Evidence from precocial species suggests that this ability might be present at the onset of life. Exploiting imprinting, an early form of learning by exposure, domestic chicks (*Gallus gallus*) successfully discriminated between XX and XY multimodal patterns [6] and between triplets of simultaneous visual elements having an AAB *vs.* ABA structure, without explicit training [7]. Newborn ducklings imprinted with visual 3D stimuli being the same or different in shape or color, at test chose novel stimuli having the same relationship as the imprinted ones, even though an alternative explanation involving stimuli visual symmetry might be in place [8].

The ability to abstract and generalize structural sequence regularities has been only marginally investigated in invertebrates, even though there are natural examples of possible stereotypical use of signal sequence production and response. For instance, the treehopper males (*Enchenopa binotata*) combine two different elements to attract females during mating: an initial whine followed by several pulses [9]. Females of this species seem to respond to the typical structure of this stimulus, being guided by a combinatory strategy [10,11]. Likewise, during the final phase of courtship, males of the black widow spider (*Latrodectus hesperus*) display an organized signal whose stereotypical structure is different from that of the signals emitted in other phases of the mating [12]. Despite this ecological evidence, very few studies have consistently investigated the ability of invertebrates to actively learn and process stimuli sequence, supported by cognitive mechanisms rather than stereotypical stimulus-response. Macquart et al. (2008) studied the ability of an ant species (*Gigantiops destructor*) to learn simple motor sequences. To reach their nest, ants were forced to explore Y-mazes according to rules of turns of increasing difficulty: from constant and repetitive turns (e.g., RRRRRRRR or LLLLLLLL; R = right, L = left) to simple alternating (e.g., RLRLRLRL), double alternating (e.g., RRLLRRLL), and irregular alternating turns (e.g., RLRRL). At test, ants correctly extended a constant-turn learning to two following novel chambers, albeit failing in applying both the simple and double alternating rule, with performance dropping from 88.8%-71.4% to 55.5%-14.3%, respectively [13]. A probable explanation implies the memorization of the maze length or number of turns needed to reach the nest, causing ants to deploy a more natural searching behavior after the first novel chamber [14]. Nevertheless, these results suggest the possible use of sensorimotor sequence learning in this species, highlighting its adaptive value with respect to costlier landmark memorization during navigation [13]. Similarly, Zhang and colleagues (1996) demonstrated the ability of honeybees to learn a sequence of flight turns to navigate a complex maze, even though their performance was poorer than learning the path by using color marks [15]. Free-flying bees presented with a serial position visual task, take into account the number and relative position of four sequential color signals to subsequently choose the rewarded left/right arm. However, while authors hypothesized a minimal role of serial pattern learning, bees seemed to rather give different salience weights depending on stimulus proximity with the reward, exhibiting a choice behavior compatible with a recency effect [16]. In this experimental design, the temporal encounter with single elements of the visual pattern was not carefully controlled, potentially leading to temporal weighting differences [16]. Overall, existing studies do not provide conclusive evidence regarding the capacity of sequence generalization in invertebrates, especially in the olfactory domain.

Nevertheless, honeybees offer a unique opportunity to try to further study their ability to form a sequence learning since they appear to be equipped with impressive cognitive and learning abilities [17]. Bees show generalization capacities, treating equally different visual stimuli based on specific physical characteristics, such as radial, circular, and bilateral symmetry [18]. Bees can also master abstract concepts such as *same/different* [19], *above/below* [20], and *right/left* [21]. They have also been tested for their ability to encode and use the temporal information of a visual stimulus to solve a subsequent task, albeit with negative results [22].

Here we aimed to investigate whether honeybees could abstract the underlying structure of a temporal sequence of three odors, and then generalize their responses to novel stimuli. We performed six experiments using the *proboscis extension response* (PER) conditioning paradigm, which allows the formation of an association between an odor stimulus and a sucrose solution that, when provided to the antennae, elicits the spontaneous extroversion of the ligula [23].

The first two experiments investigated whether bees could learn an arbitrary odor sequence, spontaneously generalize their response to novel stimuli with a similar structure but composed of novel odors, and reject different novel configurations although composed of familiar odors. We investigated the role of the position of a particular odor in the sequence and its temporal closeness to the reward, studying whether this affected bees’ memorization and recall abilities in a third control experiment. We also investigated whether differential conditioning could lead to successful discrimination of sequence structures (Exp. 4, 5). Lastly, we aimed to determine whether, by using a conditioning procedure in which a generalization strategy is favored to solve the task, a spontaneous encoding of the internal structure of odor sequences could be established and used to respond accordingly to novel stimuli (see Table 1 for a schematic representation of experimental sequence stimuli).

**Table 1:** Schematic representation of the experimental design of the six experiments.

| Experiment | Conditioning | Training Stimuli |  | Test Stimuli |  |
| --- | --- | --- | --- | --- | --- |
|  |  |  |  | Similar structure | Similar odor |
| Experiment 1 | Absolute | ABA |  | CDC | BAA |
| Experiment 2 | Absolute | CDC |  | ABA | DCC |
|  |  |  |  | First element | Second element |
| Experiment 3 | Absolute | ABA |  | AAA | BBB |
|  |  | Reinforced | Non reinforced | Congruent | Incongruent |
| Experiment 4 | Differential | ABA | BAA | CDC | DCC |
|  |  | BAA | ABA | DCC | CDC |
| Experiment 5 | Differential | ABA | AAB | CDC | CCD |
|  |  | AAB | ABA | CCD | CDC |
|  |  |  |  | Similar structure | Different structure |
| Experiment 6 | Absolute generalization | ABA, CDC |  | EFE | FEE |

## Materials and Method

All experiments were conducted at the Animal Cognition and Comparative Neuroscience Lab (ACN Lab, CIMeC, University of Trento) in Rovereto (Italy). Experiments 1, 2, and 4 were conducted from May to September 2021. Experiments 3 and 5 were conducted from April to May 2022. Experiment 6 was partly conducted in September 2021 and completed in May 2022.

### a) Subjects

Honeybee foragers (*Apis mellifera*) were obtained from colonies located in the apiary of the ACN Lab. Honeybees were anesthetized on ice until they stopped moving. Then, they were individually harnessed on metal supports. A piece of poliplak was placed on the wings in order to prevent their damage. Each fixed bee was fed with 3 µl of 30% sucrose solution (w/w) and kept in a dark, humid box for about 1 hour before starting training. In general, the day after training, all subjects were checked for their PER response following the antennal stimulation with the sucrose solution. Bees that did not exhibit the ligula extension response, were discarded from the analysis. After all experiments, subjects were released after marking them with a UNIPOSCA color to avoid cross-testing.

Overall, 324 honeybee foragers were tested (Exp.1: N = 47; Exp. 2: N = 48; Exp. 3: N = 41; Exp. 4: N = 66; Exp.5: N = 38; Exp.6: N = 84). The minimum sample size of each experiment was determined based on previous guidelines, suggesting the use of a large sample size for PER experiments (i.e., 40/50 bees per group; [24]).

### b) Apparatus and stimuli

In all experiments, the setup used to deliver the odor sequences was composed of a computerized olfactometer controlled by a MATLAB program (MATLAB R2019a, The MathWorks Inc., Natick, MA, USA), connected to six valves.

The stimuli consisted of six different odors, used according to the aim of each experiment: 3-hexanol (A), acetophenone (B), 1-nonanol (C), citral (D), benzaldehyde (E) and 2-octanone (F) (SIGMA-ALDRICH^®^), with a 1:200 concentration (5 µl odor / 1000 µl mineral oil). We selected odors that have not been proven to elicit an innate preference in honeybees [25]. For details on the sequence stimuli used see Table 1 and *Result* section.

Each odor was prepared and changed before every training session. We designed our conditioning procedure to be composed of a total of 4s of odor stimulation and 3s of sucrose delivery, with 1s overlapping [24]. This has been proposed as one of the optimal durations of stimulation in the PER paradigm [24]. At the same time, we wanted to have a nice separation between odors, to minimize the risk of odor mixture during training. Given these experimental constraints and bees’ working-memory limits of up to 3-4 elements [16], each odor sequence comprised three distinct puffs (1 second each), separated by 500 ms of air. A frontal continuous airflow and an aspirator behind the subject were used to clear the environment from the odor flow.

### c) Training procedure

All experiments followed a *proboscis extension response* conditioning procedure [24] (PER), where harnessed bees were individually conditioned to associate an odor stimulus with positive reinforcement. Each conditioning trial lasted ≈ 1 minute. The bee was placed in front of a continuous airflow for 25 seconds before the odor stimulation (4 seconds). The reinforcement (3 seconds) was delivered with a toothpick soaked in sucrose solution (i.e., ≈ 2µl of 50% sucrose solution w/w), after 3 seconds from the beginning of the odor. This allowed an overlapping of 1 second between odor and sucrose delivery. Then, after the stimulation, the bee was kept in front of the airflow for another 27 seconds and then removed. An inter-trial interval (ITI) of 10 minutes was used. The use of toothpicks does not introduce any odor bias and has been demonstrated not to be a confound for the classical CS-US learning of honeybees [26].

In experiments using the *absolute conditioning* paradigm (Exp. 1, 2, 3), bees completed a total of 10 trials, divided into 5 conditioning trials, pseudo-randomly interposed with 5 blank trials [24] (Supplementary Figure S1). In the conditioning trials, each subject was exposed to the training odor sequence, associated with the reinforcement. Conversely, during blank trials, bees were exposed to a sequence of three non-reinforced air puffs, following the identical timing sequence described above. The blank trials were used to avoid the simple learning of the puff occurrence, irrespective of the odors presented.

In experiments using the *differential conditioning* paradigm (Exp. 4, 5), bees underwent a total of 10 trials: 5 reinforced and 5 non-reinforced, pseudo-randomly presented (Supplementary Figure S2). During reinforced trials, the odor sequence was associated with positive reinforcement (i.e., 50% sucrose solution, w/w), while in non-reinforced trials, the other odor sequence was associated with water.

In experiment 6, *absolute conditioning* was used to provide a sort of generalization training. In this case, bees underwent 14 trials divided into 8 conditioned trials and 6 blank trials. During the conditioned trials, one of two odor sequences was presented (4 trials per odor sequence), while in the blank trials, bees were presented with three non-reinforced air puffs (Supplementary Figure S3). The blank trials were used to avoid the simple learning of the puff occurrence, irrespective of the odors presented. As the dependent variable for the analysis, the PER response to odor sequence was considered.

In all experiments, a spontaneous PER response at the first conditioning trial was set as the exclusion criterion.

### d) Test procedure

After training, bees underwent two testing sessions: the *memory* and the *recall* test. The test phase aimed to investigate memory formation at a different post-conditioning time. The *memory* test was conducted one hour after the end of the training, while the *recall* test was performed the morning after the training (≈ 18h delay). Both tests were conducted in probe conditions, thus not providing either reinforcement or punishment associated with the odor stimulation. An ITI of 10 minutes was set between test sequences. As the dependent variable for the analysis, the PER response to odor sequence was considered.

During the testing phase, the order of sequence presentation was randomized across subjects. Moreover, the order of sequence presentation was changed between *memory* and *recall* tests within each subject.

In experiments 1 – 3, only subjects showing the PER response to the previously trained sequence (i.e., ABA in Exp. 1 and 3, CDC in Exp. 3) during the *memory* test were considered for the analysis.

In experiments 3 and 5, we also encoded at what sequence temporal stage the PER response was observed (see Supplementary Material).

After the presentation of the last testing sequence, the presence of PER response was checked for every subject by gently stimulating its antennae with a toothpick soaked in 30% sucrose solution (w/w). Bees that did not display the response were discarded from the analysis.

### e) Statistical analyses

In the training phase, the percentage of accuracy for each trial was computed as the proportion of PER response to the stimuli sequence. The response was computed as 1 if the response was present after the odor onset, and 0 if it was not present or present before and maintained after the odor onset.

The training phase was characterized by a situation of complete separation of data, described as a condition of the allocation of all the observations in the same variable [27]. In particular, because of experiment criteria, the first trial of training was always composed of 0s (i.e., only bees not responding to the first odor presentation were considered for the analysis). Thus, the data were fitted in a *Bayesian Generalized Linear Mixed-Effect Model* (Bayesian GLMM, *bglmer* function of *blme* package) with a binomial distribution (*logit* link). The number of trials was set as a fixed factor in the model, while bee id and day were set as random factors in the model. In experiment 6, the odor type was also inserted as a fixed factor in the model. To determine the significant parameters of the models, the *anova* function (*car* package) was used and the best model was selected based on the *Akaike Information Criterion* (AIC). In GLMM, pairwise comparisons between trials were conducted using contrasts, and *p-values* were adjusted with False Discovery Rate (FDR) corrections to control for multiple comparisons (*emmeans* package) [28].

In the test phase, the percentage accuracy for each test sequence was computed as the proportion of PER response to the stimuli sequence. The response was computed as 1 if the response was present after the odor onset, and 0 if it was not present or present before and maintained after the onset of odor. The performance during the test phase was analyzed with a *Generalized Linear Mixed Effect Model* (GLMM, *glmer* function of *lme4* package) with a binomial distribution (*logit* link). The type of sequence (i.e., sequences presented during the test phase) and order of presentation of each sequence during the test were set as fixed effects in the model, while bee id and day were set as random factors. The performance during the *memory* test of experiments 1, 2, and 3, was characterized by a situation of complete separation of data (i.e., the response to the trained sequence – ABA for the first and third experiment, CDC for the second experiment – was only composed of 1s). Thus, these data were analyzed with a *Bayesian Generalized Linear Mixed-Effect Model* (Bayesian GLMM, *bglmer* function of *blme* package) with a binomial distribution (*logit* link). To determine the significant parameters of the models, the *anova* function (*car* package) was used and the best model was selected based on the *Akaike information criterion* (AIC). In GLMM, multiple comparisons between type of sequences and order of presentation were conducted using contrasts, and *p-values* were adjusted with False Discovery Rate (FDR) corrections to control for multiple comparisons (*emmeans* package) [28].

In general, when necessary models were optimized with the iterative algorithm *BOBYQA*. In all the analyses, an α-value of 0.05 was specified. All the analyses were conducted with R Studio (R, 4.1.3 version).

## 4. Results

### (a) Experiment 1 – ABA absolute conditioning paradigm

The aim of this experiment was to study whether honeybees could learn an odor sequence and then spontaneously generalize their responses to novel odor sequences having a similar structure. Honeybees were individually trained to associate the ABA odor sequences with positive reinforcement. Then, during the test phase, bees were presented with three different sequences: the trained one (i.e., ABA), a sequence having a different structure but composed of the same odorants (i.e., BAA), and a new sequence having the same structure, but composed of new odorants (i.e., CDC).

In the training phase, honeybees (n = 47) successfully learned to respond to the ABA sequence (Bayesian GLMM: PER response ∼ number of trials + (1|bee) + (1|day)). In particular, there was a significant improvement in the PER response between the first and the last training trial (post hoc analysis with FDR correction: *estimate* = −6.122, *SE* = 0.867, z*-ratio* = −7.063, *p-value* < 0.001, Fig.1a).

At *memory* test, a significant difference between sequences was found (Bayesian GLMM: PER response ∼ type of sequence + (1|bee) + (1|day); Post hoc analysis with FDR correction: ABA *vs.* BAA: *estimate* = 2.07, *SE* = 0.958, z*-ratio* = 2.16, *p-value* = 0.0308; ABA *vs.* CDC: *estimate* = 3.69, *SE* = 0.924, z*-ratio* = 3.99, *p-value* < 0.001; BAA *vs.* CDC: *estimate* = 1.62, *SE* = 0.505, z*-ratio* = 3.212, *p-value* = 0.002; Fig. 1b). Equally, significant differences among the three tested sequences were found at *recall* test (GLMM: PER response ∼ type of sequence + (1|bee) + (1|day); Post hoc analysis with FDR correction: ABA *vs.* BAA: *estimate* = −9.1, *SE* = 2.02, z*-ratio* = −4.497, *p-value* < 0.001; ABA *vs.* CDC: *estimate* = 20.4, *SE* = 3.73, z*-ratio* = 5.48, *p-value* < 0.001; BAA *vs.* CDC: *estimate* = 29.5, *SE* = 4.98, z*-ratio* = 5.935, *p-value* < 0.001; Fig. 1c).

**Figure 1:**
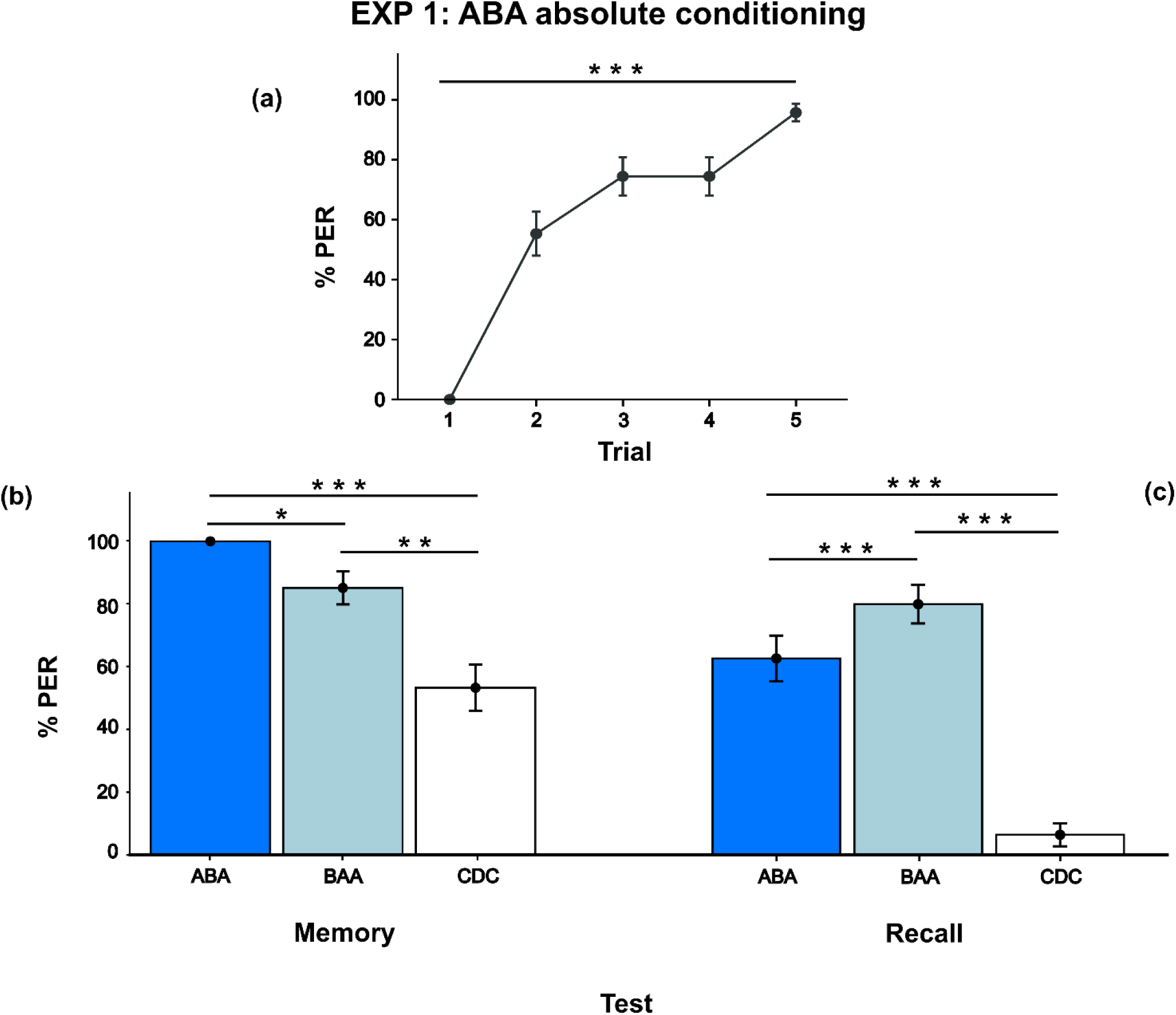
ABA absolute conditioning. Experiment 1 learning and test performance (N = 47). *(a)* Learning curve of bees trained to show the PER response when the ABA sequence (i.e., 3-hexanol – acetophenone – 3-hexanol) was presented. Bees increased their percentage of response from the first to the last trial. *(b)* Performance during the *Memory test*. *(c)* Performance during the *Recall test*. Data shown are means ± s.e.m. \**p <* 0.05, \*\**p* < 0.01, \*\*\**p* < 0.001.

Bees were thus able to learn and memorize the trained sequence (ABA), albeit without being able to generalize their response to novel odors presented with the same structure (CDC). Interestingly, at *recall* test bees showed an increased response to the novel BAA sequence (78,7%) compared to the familiar ABA one (61,7%).

### (b) Experiment 2 – CDC absolute conditioning paradigm

This experiment was a control replica of Experiment 1, to confirm the results previously obtained using a different set of odors. Honeybees were individually trained to associate the CDC odor sequence with positive reinforcement. Then, during the test phase, bees were presented with three different sequences: the trained one (i.e., CDC), a sequence having a different structure but composed of the same odorants (i.e., DCC) and a new sequence having the same structure but composed of new odorants (i.e., ABA).

In the training phase, honeybees (n = 48) successfully learned to respond to the CDC sequence (Bayesian GLMM: PER response ∼ number of trials + (1|bee) + (1|day)), with a significant improvement in the PER response between the first and last trial of training (Post hoc analysis with FDR correction: Trial 1 *vs.* Trial 5: *estimate* = −4.901, *SE* = 0.702, z*-ratio* = −6.982, *p-value* < 0.001, Fig.2a).

At *memory* test, a significant difference between CDC and ABA sequences (Bayesian GLMM: PER response ∼ type of sequence + (1|bee) + (1|day); Post hoc analysis with FDR correction: *estimate* = 4.09, *SE* = 1.07, z*-ratio* = 3.838, *p-value* < 0.001; Fig. 2b) and between DCC and ABA sequences was found (Post hoc analysis with FDR correction: *estimate* = 3.825, *SE* = 1.08, z*-ratio* = 3.554, *p-value* < 0.001; Fig. 2b). Conversely, no differences were found between the response to the CDC and DCC sequences (post hoc analysis with FDR correction: *estimate* = 0.265, *SE* = 1.27, z*-ratio* = 0.208, *p-value* = 0.835, Fig. 2b).

**Figure 2:**
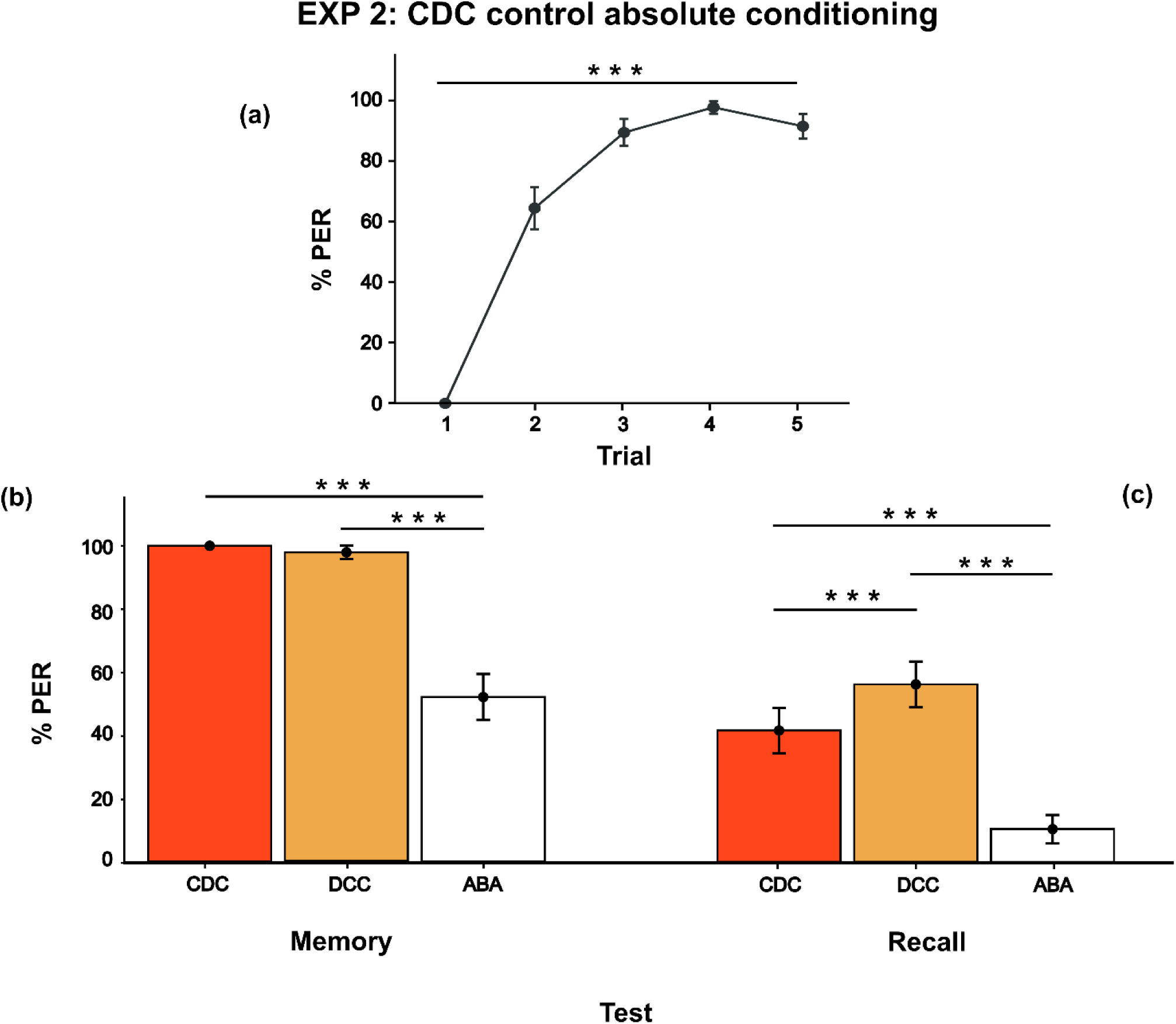
CDC absolute conditioning. Experiment 2 learning and test performance (N =48). *(a)* Learning curve of bees trained to show the PER response when the CDC sequence (i.e., 1-nonanol – citral – 1-nonanol) was presented. Bees increased their percentage of response from the first to the last trial. *(b)* Performance during the *Memory test*. *(c)* Performance during the *Recall test*. Data shown are means ± s.e.m. \*\**p* < 0.01, \*\*\**p* < 0.001.

At *recall* test, significant differences between sequences were found (GLMM: PER response ∼ type of sequence + (1|bee) + (1|day); Post hoc analysis with FDR correction: CDC *vs.* DCC: *estimate* = −14.9, *SE* = 2.93, z*-ratio* = −5.076, *p-value* < 0.001; CDC *vs.* ABA: *estimate* = 16.2, *SE* = 4.23, z*-ratio* = 3.833, *p-value* < 0.001; DCC *vs.* ABA: *estimate* = 31.1, *SE* = 6.01, z*-ratio* = 5.181, *p-value* < 0.001; Fig. 2c).

The results confirmed a lack of spontaneous generalization between the trained and novel sequences despite having the same structure (i.e., CDC and ABA). Again, at *recall* test, we observed an enhanced response to the novel DCC sequence (56,3%) with respect to the familiar CDC one (41,7%).

### (c) Experiment 3 – ABA control absolute conditioning paradigm

In experiment 3, we investigated whether some elements of the ABA sequence were memorized better thanks to their internal position in the sequence and temporal closeness to the reward. Thus, we trained honeybees to learn the association between the ABA sequence and positive reinforcement. Then, during the test phase, bees were presented with three different sequences: the trained one (i.e., ABA), a sequence composed of the A odor only (i.e., AAA), and a sequence composed of the B odor only (i.e., BBB).

Results showed that bees (n = 41) were successfully able to perform the task, by significantly increasing their response to the ABA sequence during training (Bayesian GLMM: PER response ∼ number of trials + (1|bee) + (1|day); Post hoc analysis with FDR correction: Trial 1 *vs.* Trial 5: *estimate* = −4.651, *SE* = 0.754, z*-ratio* = −6.171, *p-value* < 0.001, Fig. 3a).

**Figure 3:**
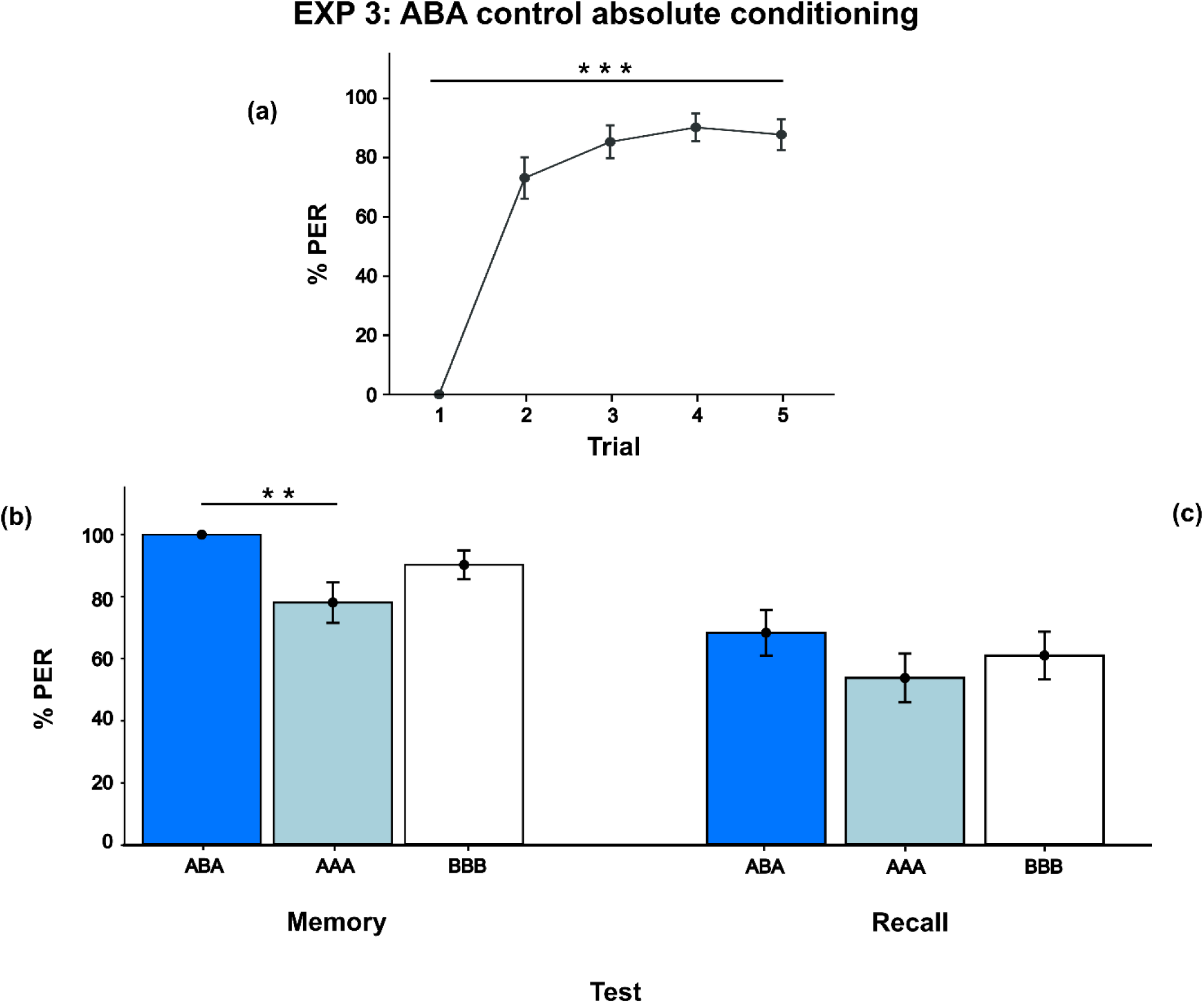
ABA control absolute conditioning. Experiment 3 learning and test performance (N =41). *(a)* Learning curve of bees trained to show the PER response when the ABA sequence (i.e., 3-hexanol – acetophenone – 3-hexanol) was presented. Bees increased their percentage of response from the first to the last trial. *(b)* Performance during the *Memory test*. *(c)* Performance during the *Recall test*. Data shown are means ± s.e.m. \**p <* 0.05.

At *memory* test, honeybees responded significantly less to the AAA sequence with respect to the ABA sequence (Bayesian GLMM: PER response ∼ type of sequence + (1|bee) + (1|day); Post hoc analysis with FDR correction, ABA *vs.* AAA: *estimate* = 2.668, *SE* = 1.029, z*-ratio* = 2.594, *p-value* = 0.029; Fig. 3b) but not to the BBB sequence (Post hoc analysis with FDR correction, ABA *vs.* BBB: *estimate* = 1.697, *SE* = 1.075, z*-ratio* = 1.579, *p-value* = 0.136; Fig. 3b). No significant differences between the novel sequences were found (Post hoc analysis with FDR correction, AAA *vs.* BBB: *estimate* = −0.972, *SE* = 0.653, z*-ratio* = −1.488, *p-value* = 0.136, Fig. 3b).

Analysis of the *recall* test showed no significant differences between models, and the simplest model was selected based on AIC value to describe the data (GLMM: PER response ∼ 1 + (1|id); Fig. 3c). This highlighted a non-significant effect of the type of sequence as a factor.

We investigated whether the results previously obtained at *recall* test of experiments 1 and 2 (i.e., higher response to novel unfamiliar sequences composed of the same odors with respect to the trained familiar ones) could be explained by a different encoding of one element of the sequence (i.e., odor A or odor B) due to their position in the sequence and closeness to the reward delivery. The results of *recall* test did not support this hypothesis since a similar response to the A and B odors was found.

### (d) Experiment 4 – ABA *vs.* BAA differential conditioning paradigm

The purpose of Experiment 4 was to determine whether the specific learning of the structure of an odor sequence could be obtained by exploiting a differential conditioning paradigm. Two independent groups of honeybees (group 1: n = 36; group 2: n = 30) were trained to discriminate between the ABA and BAA sequences (first group: ABA reinforced, BAA non-reinforced; second group: BAA reinforced and ABA non-reinforced). Then, at test, their response to the previously reinforced and non-reinforced sequences was scored, together with their spontaneous PER response to congruent (i.e., CDC for the first group; DCC for the second group) and incongruent (i.e., DCC for the first group; CDC for the second group) sequences composed of novel odors.

Results showed a significant effect on the number of trials, with a positive increase from the first to the last trial (Bayesian GLMM: PER response ∼ number of trials + type of sequence + number of trials: type of sequence + (1|bee) + (1|day); Post hoc analysis with FDR correction: Trial 1 *vs.* Trial 5: *estimate* = −5.998, *SE* = 0.574, z*-ratio* = −10.446, *p-value* < 0.001). However, no differences between sequences were found (Post hoc analysis with FDR correction, ABA reinforced *vs.* BAA non-reinforced: *estimate* = −0.339, *SE* = 0.387, z*-ratio* = −0.876, *p-value* = 0.536; ABA reinforced *vs.* BAA reinforced: *estimate* = 0.236, *SE* = 0.722, z*-ratio* = 0.326, *p-value* = 0.744; ABA reinforced *vs.* ABA non-reinforced: *estimate* = 1.025, *SE* = 0.728, z*-ratio* = 1.409, *p-value* = 0.318; ABA non-reinforced *vs.* BAA reinforced: *estimate* = 0.79, *SE* = 0.36, z*-ratio* = 2.193, *p-value* = 0.169; ABA non-reinforced *vs.* BAA non-reinforced: *estimate* = −1.364, *SE* = 0.762, z*-ratio* = - 1.79, *p-value* = 0.22; BAA reinforced *vs.* BAA non-reinforced: *estimate* = −0.575, *SE* = 0.755, z*-ratio* = −0.761, *p-value* = 0.536). The interaction term showed a significant impact on data. However, when we analyzed the crucial comparisons (i.e., between reinforced and non-reinforced sequences presented to the same group and at a given trial), post-hoc analyses did not reveal significant differences, confirming the incapacity of bees to discriminate between sequences during the training phase.

Thus, we collapsed the two experimental groups, creating two new sequence categories (i.e., reinforced, and non-reinforced sequences), and analyzed the data. The model (Bayesian GLMM: PER response ∼ number of trials + type of sequence + number of trials:type of sequence + (1|bee) + (1|day)) highlighted a significant impact of the interaction factor. However, when the crucial comparisons were analyzed (i.e., comparison between reinforced and non-reinforced sequences at a given trial), only a significant difference between sequences at the first trial was present (Post hoc analysis with FDR correction: *estimate* = 2.004, *SE* = 0.713, z*-ratio* = 2.813, *p-value* = 0.009; Fig. 4a). Irrespective of the type of sequence, an improvement from the first to the last trial was observed (Post hoc analysis with FDR correction: *estimate* = −6.05, *SE* = 0.589, z*-ratio* = - 10.265, *p-value* < 0.001; Fig. 4a). There was no difference in PER response to reinforced and non-reinforced sequences (Post hoc analysis with FDR correction: *estimate* = 0.33, *SE* = 0.264, z*-ratio* = 1.25, *p-value* = 0.211).

**Figure 4:**
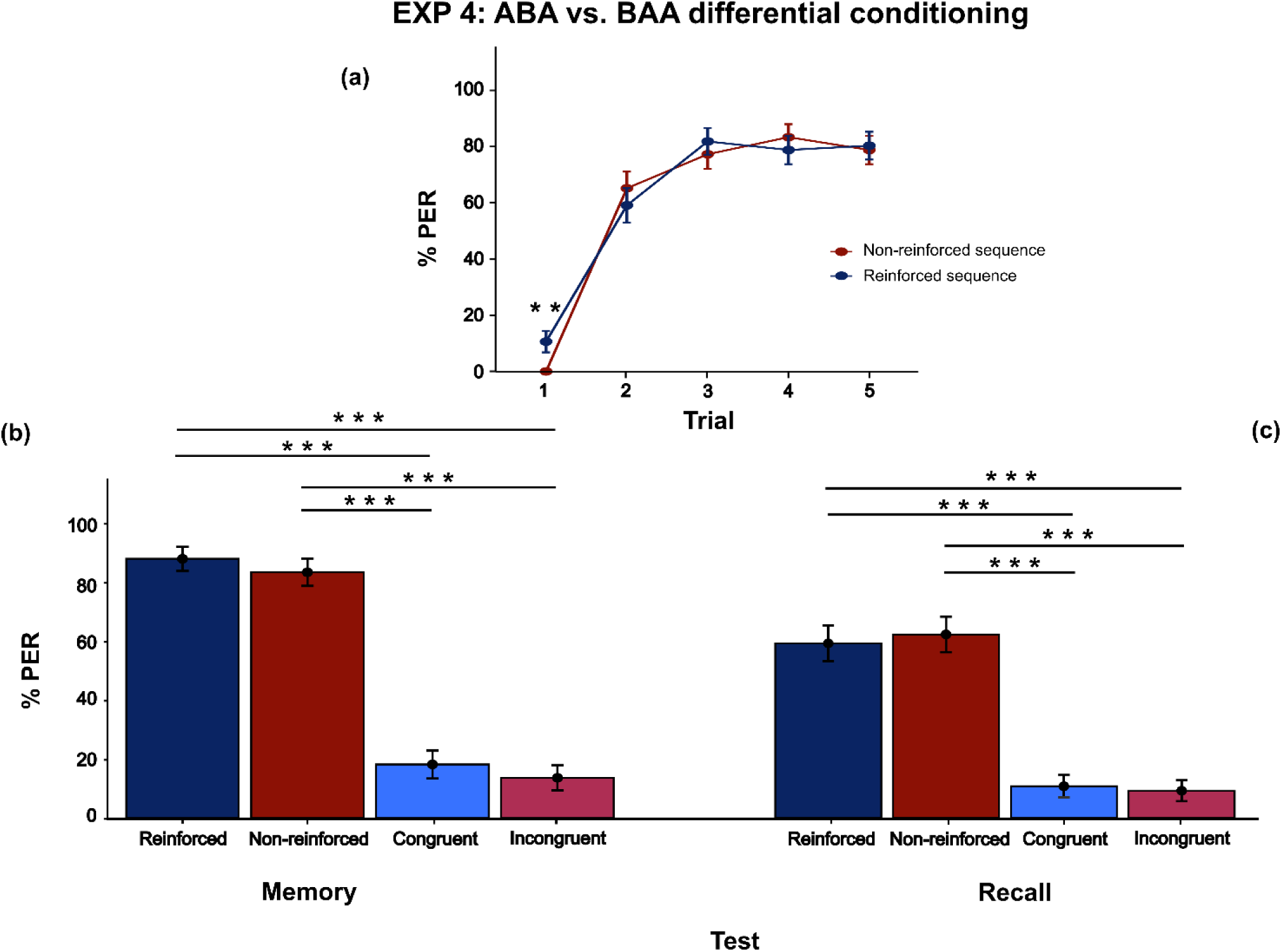
ABA *vs.* BAA differential conditioning. Experiment 4 learning and test performance (N = 66). *(a)* Learning curve of bees trained to discriminate between the ABA (i.e., 3-hexanol – acetophenone – 3-hexanol) and BAA (i.e., acetophenone – 3-hexanol – 3-hexanol) sequences. *Reinforced* and *non-reinforced* sequences were defined as sequences associated during training with sucrose solution and water, respectively. *Congruent* and *incongruent* sequences were defined as sequences composed of novel odors having the same structure as previously reinforced and non-reinforced sequences, respectively. *(b)* Performance during the *Memory test*. *(c)* Performance during the *Recall test*. Data shown are means ± s.e.m. \*\**p* < 0.01, \*\*\**p* < 0.001.

During the test phase, four different sequences were presented in probe conditions: previously reinforced (i.e., ABA for group 1, BAA for group 2), previously non-reinforced (i.e., BAA for group 1, ABA for group 2), congruent (i.e., sequence with new odor and same structure of the previously reinforced one: CDC for group 1, DCC for group 2) and incongruent (i.e., sequence with new odor and same structure of the previously non-reinforced one: DCC for group 1 and CDC for group 2).

At *memory* test, no differences were found between the reinforced and non-reinforced sequences, (GLMM: PER response ∼ type of sequence + (1|bee) + (1|day); Post hoc analysis with FDR correction: *estimate* = 1.24, *SE* = 0.994, z*-ratio* = 1.247, *p-value* = 0.234; Fig. 4b). The response to the congruent and incongruent sequences was not significantly different (Post hoc analysis with FDR correction: *estimate* = 1.08, *SE* = 0.905, z*-ratio* = 1.189, *p-value* = 0.234; Fig. 4b). Bees showed a significantly lower PER response to both congruent and incongruent sequences with respect to the reinforced and non-reinforced ones (Post hoc analysis with FDR correction: reinforced *vs.* congruent: *estimate* = 12.92, *SE* = 2.231, z*-ratio* = 5.791, *p-value* < 0.001; reinforced *vs.* incongruent: *estimate* = 13.99, *SE* = 2.463, z*-ratio* = 5.681, *p-value* < 0.001; non-reinforced *vs.* congruent: *estimate* = 11.68, *SE* = 1.913, z*-ratio* = 6.104, *p-value* < 0.001; non-reinforced *vs.* incongruent: *estimate* = 12.75, *SE* = 2.159, z*-ratio* = 5.909, *p-value* < 0.001; Fig. 4b).

Similar results were found at *recall* test, where bees did not differentiate between the reinforced and non-reinforced sequences (Post hoc analysis with FDR correction: *estimate* = −0.353, *SE* = 0.598, z*-ratio* = −0.591, *p-value* = 0.637; Fig. 4c) and between the congruent and incongruent sequences (Post hoc analysis with FDR correction: *estimate* = 0.456, *SE* = 0.966, z*-ratio* = 0.472, *p-value* = 0.637; Fig. 4c). Bees showed a significantly lower PER response to both congruent and incongruent sequences with respect to the reinforced and non-reinforced ones (Post hoc analysis with FDR correction: reinforced *vs.* congruent: *estimate* = 7.283, *SE* = 1.662, z*-ratio* = 4.383, *p-value* < 0.001; reinforced *vs.* incongruent: *estimate* = 7.739, *SE* = 1.788, z*-ratio* = 4.328, *p-value* < 0.001; non-reinforced *vs.* congruent: *estimate* = 7.637, *SE* = 1.713, z*-ratio* = 4.458, *p-value* < 0.001; non-reinforced *vs.* incongruent: *estimate* = 8.093, *SE* = 1.839, z*-ratio* = 4.402, *p-value* < 0.001; Fig. 4c).

In summary, in experiment 4 honeybees neither discriminated between ABA and BAA sequences nor generalized their response to novel congruent (i.e., having the same structure as the previously reinforced one) or incongruent (i.e., having the same structure as the previously non-reinforced sequence) sequences, highlighting an inability to discriminate olfactory stimuli based on their internal structure. This could be a result of bees mainly processing the last sequence element that, in this case, was the same between sequences (i.e., A odor in the last position).

### (e) Experiment 5 – ABA *vs.* AAB differential conditioning paradigm

In this experiment, we aim to investigate whether the last element of the sequence was more informative and thus used by bees to discriminate between two odor structures. We trained two independent groups of honeybees (group 1: n = 20; group 2: n = 18) to discriminate between the ABA and AAB sequences. The procedure was the same as described for experiment 4. We also encoded the occurrence of the PER response in relation to the presentation of a specific element of the sequence (i.e., first, second or third element).

At training, results showed a significant effect of the number of trials, with an increase in response from the first to the last trial (Bayesian GLMM: PER response ∼ number of trials + type of sequence + (1|bee) + (1|day); Post hoc analysis with FDR correction: *estimate* = −4.817, *SE* = 0.576, z*-ratio* = −8.363, *p-value* < 0.001). The model highlighted a difference between the two reinforced sequences (Post hoc analysis with FDR correction: *estimate* = 1.372, *SE* = 0.511, z*-ratio* = 2.686, *p-value* = 0.043). However, no other differences among sequences were found (Post hoc analysis with FDR correction: ABA reinforced *vs.* ABA non-reinforced: *estimate* = 0.584, *SE* = 0.515, z*-ratio* = 1.132, *p-value* = 0.309; ABA reinforced *vs.* AAB non-reinforced: *estimate* = 0.763, *SE* = 0.399, z*-ratio* = 1.911, *p-value* = 0.112; AAB reinforced *vs.* ABA non-reinforced: *estimate* = −0.788, *SE* = 0.403, z*-ratio* = −1.955, *p-value* = 0.112; AAB reinforced *vs.* AAB non-reinforced: *estimate* = −0.61, *SE* = 0.499, z*-ratio* = −1.222, *p-value* = 0.309; ABA non-reinforced *vs.* AAB non-reinforced: *estimate* = 0.179, *SE* = 0.509, z*-ratio* = 0.351, *p-value* = 0.725).

The significant difference between the two rewarded sequences was not considered a reliable indicator of differences in the ability of bees to better learn one of the sequences over the other. Honeybees of both groups could not discriminate between rewarded and non-rewarded sequences during training. Thus, we decided to proceed with the analysis of the totality of data, collapsing the two groups together.

Thus, we created two new sequence categories, as in experiment 4 (i.e., reinforced, and non-reinforced sequences), and analyzed the data. The number of trials had a significant impact on the bees’ performance, with an improvement from the first to the last trial (Bayesian GLMM: PER response ∼ number of trials + (1|bee) + (1|day); Post hoc analysis with FDR correction: *estimate* = −4.69, *SE* = 0.564, z*-ratio* = −8.314, *p-value* < 0.001; Fig. 5a). The model did not report any significant impact of the sequence factor (i.e., reinforced *vs.* non-reinforced) on the data.

**Figure 5:**
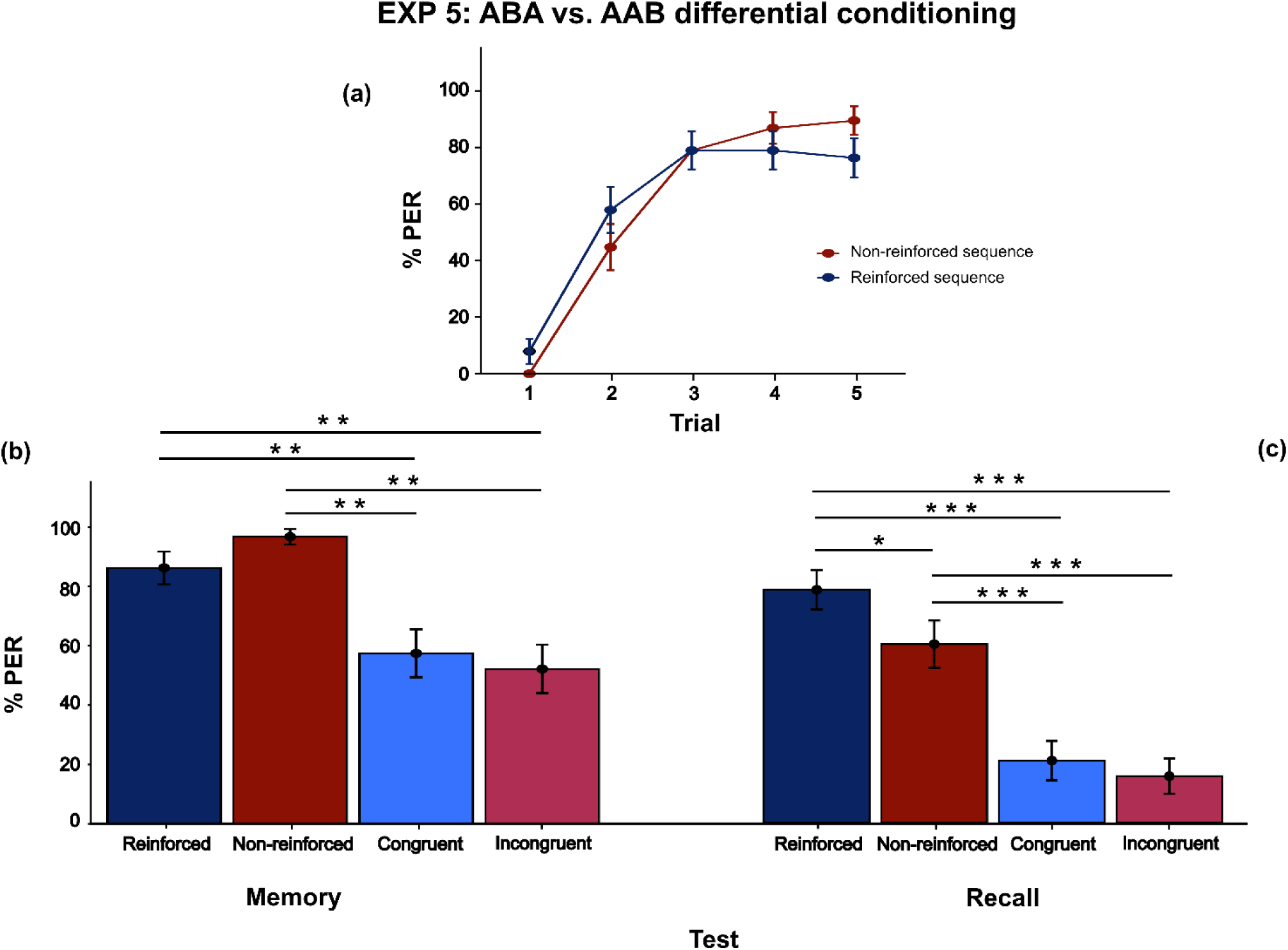
ABA *vs.* AAB differential conditioning. Experiment 5 learning and test performance (N = 38). *(a)* Learning curve of bees trained to discriminate between the ABA (i.e., 3-hexanol – acetophenone – 3-hexanol) and AAB (i.e., 3-hexanol – 3-hexanol - acetophenone) sequences. *Reinforced* and *non- reinforced* sequences were defined as sequences associated during training with sucrose solution and water, respectively. *Congruent* and *incongruent* sequences were defined as sequences composed of novel odors having the same structure as previously reinforced and non-reinforced sequences, respectively. *(b)* Performance during the *Memory test*. *(c)* Performance during the *Recall test*. Data shown are means ± s.e.m. \**p <* 0.05, \*\**p* < 0.01, \*\*\**p* < 0.001.

During the test phase, we presented the four different sequences in probe conditions: previously reinforced (i.e., ABA for group 1, AAB for group 2), previously non-reinforced (i.e., AAB for group 1, ABA for group 2), congruent (i.e., sequence with new odor and same structure of the previously reinforced one: CDC for group 1, CCD for group 2) and incongruent (i.e., sequence with new odor and same structure of the previously non-reinforced one: CCD for group 1 and CDC for group 2).

At *memory* test, no differences were found neither between the reinforced and non-reinforced sequences (GLMM: PER response ∼ type of sequence + (1|bee) + (1|day); Post hoc analysis with FDR correction: *estimate* = −1.85, *SE* = 1.157, z*-ratio* = −1.60, *p-value* = 0.132; Fig. 5b) nor between congruent and incongruent sequences (Post hoc analysis with FDR correction: *estimate* = 0.273, *SE* = 0.524, z*-ratio* = 0.52, *p-value* = 0.603; Fig. 5b). Conversely, bees responded significantly less to congruent and incongruent sequences with respect to the reinforced and non-reinforced ones (Post hoc analysis with FDR correction: reinforced *vs.* congruent: *estimate* = 1.895, *SE* = 0.675, z*-ratio* = 2.808, *p-value* = 0.008; reinforced *vs.* incongruent: *estimate* = 2.168, *SE* = 0.689, z*-ratio* = 3.147, *p-value* = 0.003; non-reinforced *vs.* congruent: *estimate* = 3.745, *SE* = 1.153, z*-ratio* = 3.247, *p-value* = 0.003; non-reinforced *vs.* incongruent: *estimate* = 4.02, *SE* = 1.166, z*-ratio* = 3.446, *p-value* = 0.003; Fig. 5b).

At *recall* test, bees successfully discriminate between reinforced and non-reinforced sequences (GLMM: PER response ∼ type of sequence + (1|bee) + (1|day); Post hoc analysis with FDR correction: *estimate* = 1.571, *SE* = 0.738, z*-ratio* = 2.128, *p-value* = 0.04; Fig. 5c). Bees responded significantly less to congruent and incongruent sequences with respect to the reinforced and non-reinforced ones (Post hoc analysis with FDR correction: reinforced *vs.* congruent: *estimate* = 4.683, *SE* = 1.086, z*-ratio* = 4.314, *p-value* < 0.001; reinforced *vs.* incongruent: *estimate* = 5.236, *SE* = 1.166, z*-ratio* = 4.491, *p-value* < 0.001; non-reinforced *vs.* congruent: *estimate* = 3.112, *SE* = 0.865, z*-ratio* = 3.596, *p-value* < 0.001; non-reinforced *vs.* incongruent: *estimate* = 3.665, *SE* = 0.945, z*-ratio* = 3.880, *p-value* < 0.001; Fig. 5c). Again, no differences in the PER response to congruent and incongruent sequences emerged (Post hoc analysis with FDR correction: *estimate* = 0.553, *SE* = 0.753, z*-ratio* = 0.735, *p-value* = 0.462; Fig. 5c).

As in experiment 4, honeybees trained to discriminate between ABA and AAB sequences did not show a spontaneous generalization of response to novel congruent and incongruent sequences (i.e., having the same structure of previously reinforced and non-reinforced sequences, respectively). Bees confirmed the inability to discriminate between ABA and AAB sequences during the training and *memory* test phase. However, this ability emerged during the *recall* test, suggesting the use of different strategies based on the position of particular sequence elements in long-term memory formation (see Discussion).

### (f) Experiment 6 – Generalization conditioning paradigm

In this experiment, a group of honeybees (n = 84) was trained to learn to respond to two sequences having the same structure but different odors (i.e., ABA and CDC). Then, the ability of bees to generalize to a new odor sequence having the same internal structure (i.e., EFE) and to differentiate it from a sequence having a different structure (i.e., FEE) was tested.

During the training phase, the best model highlighted a significant effect of the number of trials on the percentage of response, with PER accuracy increasing from the first to the last trial (Bayesian GLMM: PER response ∼ number of trials + (1|bee) + (1|day); Post hoc analysis with FDR correction: *estimate* = −5.072, *SE* = 0.554, z*-ratio* = −9.152, *p-value* < 0.001; Fig. 6a). The type of odor presented (i.e., ABA - 3-hexanol/acetophenone/3- hexanol - and CDC - 1-nonanol/citral/1-nonanol) was not highlighted as a significant factor by model, suggesting that during training, honeybees responded equally to the two sequences.

**Figure 6:**
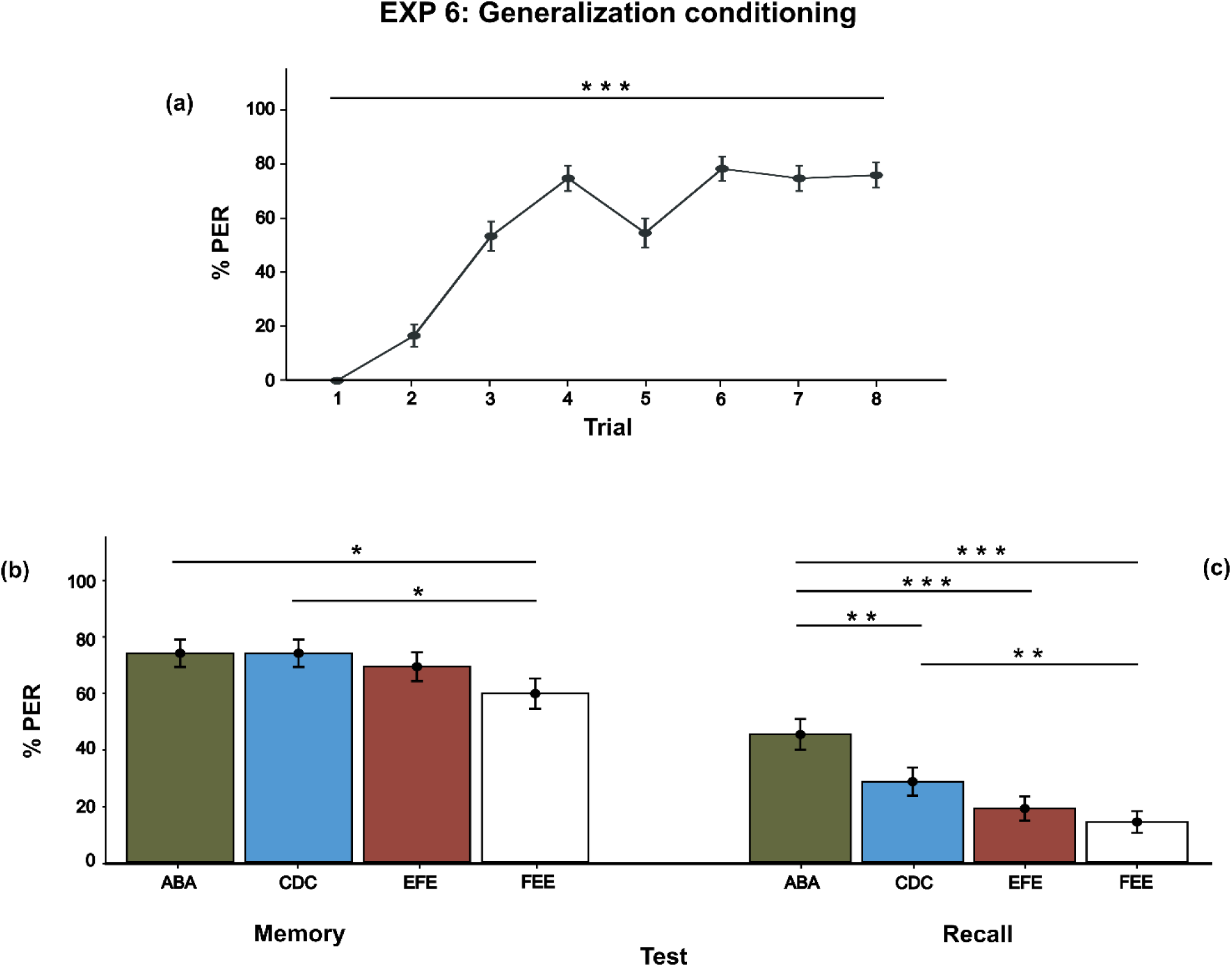
Generalization conditioning. Experiment 6 learning and test performance (N = 84). *(a)* Learning curve of bees trained to show the PER response when the ABA and CDC sequences (i.e., ABA: 3-hexanol – acetophenone – 3-hexanol; CDC: 1-nonanol – citral – 1-nonanol) were presented. Bees increased their percentage of response from the first to the last trial, without any differences between the proportion of response to the two sequences. *(b)* Performance during the *Memory test*. *(c)* Performance during the *Recall test*. Data shown are means ± s.e.m. \**p <* 0.05, \*\**p* < 0.01, \*\*\**p* < 0.001.

At *memory* test, honeybees responded significantly less to the FEE sequence with respect to the trained ones (GLMM: PER response ∼ type of sequence + (1|bee) + (1|day); Post hoc analysis with FDR correction: ABA *vs.* FEE: *estimate* = 1.082, *SE* = 0.44, z*-ratio* = 2.461, *p-value* = 0.042; CDC *vs.* FEE: *estimate* = 1.082, *SE* = 0.44, z*-ratio* = 2.461, *p-value* = 0.042; Fig. 6b), but not to the new sequence having the same structure (Post hoc analysis with FDR correction: ABA *vs.* EFE: *estimate* = 0.384, *SE* = 0.44, z*-ratio* = 0.872, *p-value* = 0.459; CDC *vs.* EFE: *estimate* = 0.384, *SE* = 0.44, z*-ratio* = 0.872, *p-value* = 0.459; Fig. 6b). No differences were found between the EFE and FEE sequences during the test (Post hoc analysis with FDR correction: *estimate* = 0.699, *SE* = 0.424, z*-ratio* = 1.648, *p-value* = 0.199; Fig. 6b).

Results of *recall* test showed a significant effect of type of sequence (GLMM: PER response ∼ type of sequence + order of presentation + (1|bee) + (1|day); Post hoc analysis with FDR correction: ABA *vs.* CDC: *estimate* = 1.252, *SE* = 0.444, z*-ratio* = 2.818, *p-value* = 0.009; ABA *vs.* EFE: *estimate* = 2.126, *SE* = 0.502, z*-ratio* = 4.232, *p-value* < 0.001; ABA *vs.* FEE: *estimate* = 2.677, *SE* = 0.554, z*-ratio* = 4.828, *p-value* < 0.001; CDC *vs.* EFE: *estimate* = 0.874, *SE* = 0.48, z*-ratio* = 1.822, *p-value* = 0.082; CDC *vs.* FEE: *estimate* = 1.425, *SE* = 0.522, z*-ratio* = 2.732, *p-value* = 0.009; EFE *vs.* FEE: *estimate* = 0.551, *SE* = 0.531, z*-ratio* = 1.038, *p-value* = 0.299; Fig. 6c). Post hoc analysis did not showed a significant difference between presentation order (first *vs.* second: *estimate* = - 0.507, *SE* = 0.525, z*-ratio* = −0.966, *p-value* = 0.401; first *vs.* third: *estimate* = −1.138, *SE* = 0.55, z*-ratio* = −2.068, *p-value* = 0.116; first *vs.* fourth: *estimate* = −1.316, *SE* = 0.522, z*- ratio* = −2.522, *p-value* = 0.07; second *vs.* third: *estimate* = −0.63, *SE* = 0.496, z*-ratio* = - 1.271, *p-value* = 0.306; second *vs.* fourth: *estimate* = −0.808, *SE* = 0.485, z*-ratio* = −1.666, *p-value* = 0.191; third *vs.* fourth: *estimate* = −0.178, *SE* = 0.483, z*-ratio* = −0.369, *p-value* = 0.712).

Results of *memory* test showed a decreased response for the sequence composed of novel odors presented with a novel structure (i.e., FEE) compared to sequences having a familiar structure, irrespective of the constituent odors (i.e., ABA, CDC, EFE). At *recall* test, this result was not replicated as a generally higher response to ABA and CDC sequence with respect to the novel ones (i.e., EFE, FEE) was found.

## Discussion

We systematically investigated whether honeybees could extract an underlying temporal odor regularity from the environment. To this aim, we adapted the classical PER paradigm and observed bees’ ability to extrapolate an underlying structure of consecutively presented odors forming a sequence.

Experiments 1 and 2 revealed that bees associated the trained sequences with positive reinforcement (Fig. 1a, 2a) without, however, being spontaneously able to generalize this response to novel odors presented with the same structure (i.e., CDC, Exp. 1, Fig. 1b, 1c; ABA, Exp. 2, Fig. 2b, 2c). In these conditions, honeybees’ behavior suggested a primary encoding of the individual odor properties instead of the entire sequence order presentation. Interestingly, bees also showed a significant increase in response to novel-structured sequences at *recall* test (i.e., BAA, Exp. 1, Fig. 1c; DCC, Exp. 2, Fig. 2c). This outcome was not driven by a different retrieval of one of the two odors constituting the sequence stimulus (see Results of Exp.3, *recall* test). When we analyzed the exact moment of PER display (Exp. 3), bees preferentially responded to both the A and B odors presented in the first position, suggesting again a similar encoding of the two odors irrespective of their ordering structure and temporal closeness to the reward (Supplementary Figure S4). The existence of a better memorization of the peculiar dyad composed of second and third elements (e.g., BA, DC) could be another possible explanation for this result. This option might have been favored by the stricter temporal contingency of the second and third elements with the sucrose solution provided as reinforcement (i.e., just before and during the sucrose delivery, respectively).

It might be argued that bees could have learned an A+B odor mixture compound, instead of a sequential ABA odor presentation where each elemental component has a unique position in the sequence. First and foremost, we temporally separate the three sequential odor puffs, providing a continuous airflow to clean the environment and minimize any possible odor overlapping. Furthermore, results from the first two experiments might help to answer this concern. If bees were mainly learning to respond to a compound formed by mixed odors (e.g., A+B rather than ABA in Exp. 1; C+D rather than CDC in Exp. 2), we would not have expected to observe the significantly different proportion of response between trained sequences (i.e., ABA, Exp. 1; CDC, Exp.2) and novel structure sequences composed of familiar odors in the *recall* test of Experiment 1 and 2 (i.e., BAA, Exp. 1; DCC, Exp. 2) as they equally carried the ‘mixture’ cue, despite the temporal order of their composing elements.

The use of differential conditioning highlighted again an inability of bees to actively differentiate between ABA and BAA (Exp. 4; Fig. 4), or ABA and AAB structures (Exp. 5; Fig. 5). Furthermore, the significant difference in the proportion of response between ABA and AAB sequences at the *recall* test (Fig. 5c), could not rule out a discrimination capacity based on the last element dissimilarity, rather than a complete sequence learning.

Observation of the exact moment of PER display (Exp. 5: Supplementary Figure S5) might suggest the presence of an additional *response to the odor change* strategy. When novel odors were presented (i.e., CDC, CCD; *recall* test of Exp. 5), a considerable number of bees responded to the D odor, namely the odor different from the first one presented (C; Supplementary Figure S5). Similarly, when the trained ABA and AAB stimuli were presented, bees were more likely to respond not only to the first odor (A) but also to the subsequent different odor (B) irrespective of the position of the latter (either second or third in the sequence; Supplementary Figure S5), supporting a possible spontaneous use of this strategy. Arguably, a generalization response among citral (D) and 3-hexanol (A) or between citral (D) and acetophenone (B) could explain these results. However, past research demonstrated bees’ discrimination ability between citral and 1-hexanol compounds [29]. Since a strong generalization at both behavioral [30] and neuronal levels between different hexanols (i.e., 1-hexanol, 2-hexanol, 3-hexanol; [31]) has been suggested, we could acknowledge a low probability for bees to generalize their response between citral and 3-hexanol in our study. Likewise, citral and acetophenone have different neuronal activity, sustained by specific glomeruli activation in the bee brain [31,32]. The low generalization response between sequences in Exp. 1 and 2 (i.e., ABA *vs.* CDC) also supports the idea that a generalization among odors was unlikely to be present in our research.

Honeybees showed a response congruent with structure similarity (i.e., ABA, CDC, and EFE) and dissimilarity strategy (i.e., FEE) in experiment 6 (*memory* test, Fig. 6b). The capacity to generalize the structure across different odors was present only one hour after training. When we tested the long-term memory capacity, a higher response for the two trained sequences (i.e., ABA, CDC) was found, together with a significant difference between them (*recall* test, Fig. 6c). This latter difference might be caused by a more robust memorization of the 3-hexanol and acetophenone odors (composing the ABA stimulus) compared to 1-nonanol and citral odors (composing the CDC stimulus). Results of the *recall* test of experiments 1 and 2 support this hypothesis, with the proportion of bees responding to the ABA-trained sequence (61,7%, Exp. 1; Fig. 1c) being higher than the proportion of subjects responding to the CDC-trained sequence (41,7%, Exp. 2; Fig. 2c). We did not find a significant difference between novel stimuli with either a familiar or a non-familiar structure (i.e., EFE, FEE). This might be an effect of the relatively short training (i.e., 8-trials training providing only two odor sequences with the same structure). Future research should investigate whether a more extended and diverse training procedure could enhance the ability of bees to generalize their response and differentiate between novel odors with familiar and unfamiliar structures. In visual discrimination tasks, the use of appetitive-aversive conditioning has been shown to enhance honeybees’ solving performance [33]. Unfortunately, the use of quinine solution as the aversive stimulus is not fully supported during the PER conditioning paradigm due to its toxicity rate (just under 60% mortality for 100mM quinine concentration; [34]).

Bees showed different patterns of response at *memory* and *recall* tests in our experiments (e.g., lower rate of response during *recall* tests with respect to *memory* tests). In Pavlovian conditioning, memory retrieval can depend on several factors that can be modulated during training: interstimulus interval (i.e., time intercourse between CS and US; ISI), intertrial interval (i.e., time intercourse between subsequent trials; ITI), and the total number of trials [35]. Moreover, a process of memory decay can also explain the differences in response during *recall* tests [36].

Bees seem to use different strategies when presented with the sequence discrimination task. This should not be surprising as it has been demonstrated that the use of different conditioning procedures (i.e., absolute or differential) could determine variations in stimuli discrimination [37]. The discovery of alternative strategies implementation is pivotal for understanding honeybees’ learning processes, the construction of abstract knowledge, and which experimental tools are necessary to let it occur. Avarguès-Weber et al. (2011) demonstrated the existence of an abstract *above/below* categorization rule based on the element’s spatial relationship during honeybees’ visual discrimination task, without relying on target absolute position [20, 21]. High-speed videography analysis suggested that bees could solve the task by employing a scanning behavior of the lower visual item before making a decision [38], without necessarily relying on a more abstract conceptual rule. However, it is worth noticing that even in this latter study, evidence for above/below categorization was found [38]. After extensive training, bumblebees trained in a *small* vs. *large* visual shape discrimination, opted for a simpler win-stay/lose-switch strategy, albeit still retaining the learning of a relative size rule [39]. Despite the apparent incompatibility of the described evidence, it has been suggested that the development of different mechanisms supporting low and high-cognitively demanding behaviors could be adaptive in ecological situations [39].

The experimental design of the present study involved a presentation of odor stimuli that might appear to be as not ecologically relevant, for it could be argued that foragers might not encounter temporally separated odor bouquets during their flight, so that any direct comparison between our experimental conditions and natural behavior may appear weak. However, to investigate the ability of bees to learn and generalize among sequences, it is mandatory to use a design that allows for rigorous control over independent variables (e.g., the timing of stimuli occurrence) and presentation of ecologically salient sequences with which honeybees could create an association with positive reinforcement (i.e., odor sequences). Thus, the main goal of our experimental design was not to mimic natural behavior and study the underlying cognitive process but to provide an effective experimental method to investigate the presence of a specific cognitive ability in an insect species, shedding light on our understanding of bees’ cognition.

The capacity to extrapolate regularities from the environment is thought to be of high adaptive value as it allows animals to conform their behavior in response to stimuli that share common regular characteristics despite being composed of different elements, ultimately helping to discover the underlying environmental structure [7]. Bees are equipped with a minimal circuit favoring the construction of categorical and generalization processes rather than overloading the memory capacity [21]. Moreover, bees communicate through a sensorimotor display known as the waggle dance, which comprises elements presented according to a sequential structure [40]. Therefore, an abstraction capacity related to temporal sequence structures in bees could be hypothesized. However, our results do not provide a further and clear demonstration of the ability of bees to extrapolate temporal contingencies and successfully generalize that knowledge to novel stimuli. Rather, our findings support the implementation of simpler rules to address an odor sequence discrimination task. Still, we do not exclude the possibility that the ability to extrapolate a structure regularity might be present or better developed in other sensory modalities (e.g., tactile) in honeybees.

## Acknowledgments

We would like to thank Francesco Rughi for his help in the data collection of experiments 2, 4, and 6, and Bastien Lemaire, Mirko Zanon, and Orsola Rosa Salva for their useful comments on the previous version of this manuscript.

## Author contributions

M.B. and G.V. conceived the study. M.B. designed the experiment, analyzed data, and wrote the initial version of the manuscript. G.V. contributed to the study design, data interpretation, and editing of the manuscript.

## Ethics and Consent section

The manuscript presents research on animals that do not require ethical approval for their study.

## Data and code availability

Raw data have been deposited at the Mendeley Data repository (DOI: 10.17632/bcxx3j4y32.1).

