## Supplementary Materials for "Odor sequence learning in honeybees: insights from olfactory classical conditioning paradigm"

**SUPPLEMENTAL ITEMS**

**FIGURES**

**Supplementary Figure 1. Schematic representation of the absolute conditioning training paradigm, Related to Figures 1a, 2a, and 3a.**

During the training of Exp. 1, Exp. 2, and 3 honeybees were presented with a total of 10 trials, divided into 5 experimental trials and 5 blank trials. During the blank trials (represented as white bars) a sequence of three puffs of air was provided without any positive reinforcement. Conversely, in the experimental trials, the odor sequence (represented as blue bars) was provided to the bee in association with a drop of sucrose solution (represented as a light green bar).

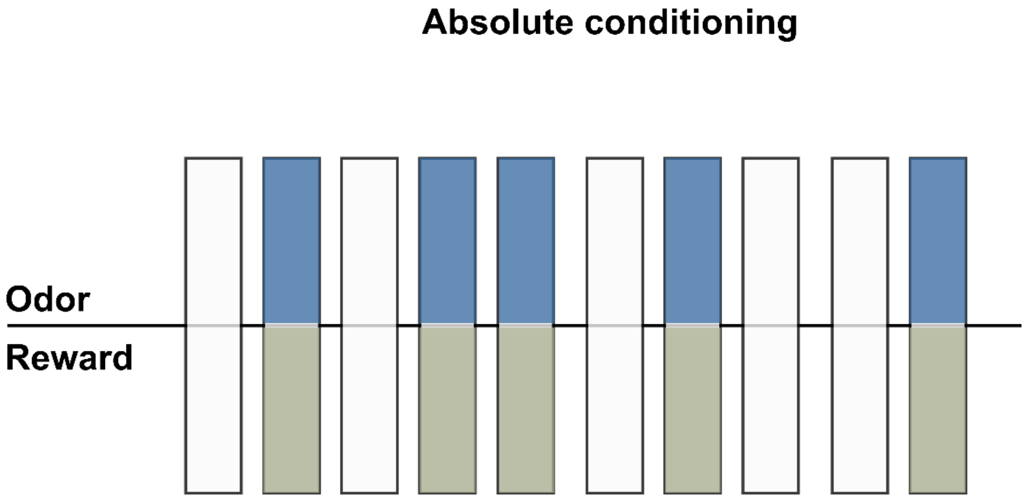

**Supplementary Figure 2. Schematic representation of the differential conditioning training paradigm, Related to Figures 4a and 5a.**

During the training of Exp. 4 and 5 honeybees were presented with a total of 10 trials, divided into 5 correct trials and 5 incorrect trials. Depending on the experimental group, during the correct trials bees were presented with a certain odor sequence (either ABA, BAA, or AAB; represented by blue bars) in association with the sucrose solution (represented as light green bars). Conversely, during the incorrect trials, a different sequence was presented (either ABA, BAA, or AAB; represented by red bars) in association with a neutral stimulus (water; represented as white bars).

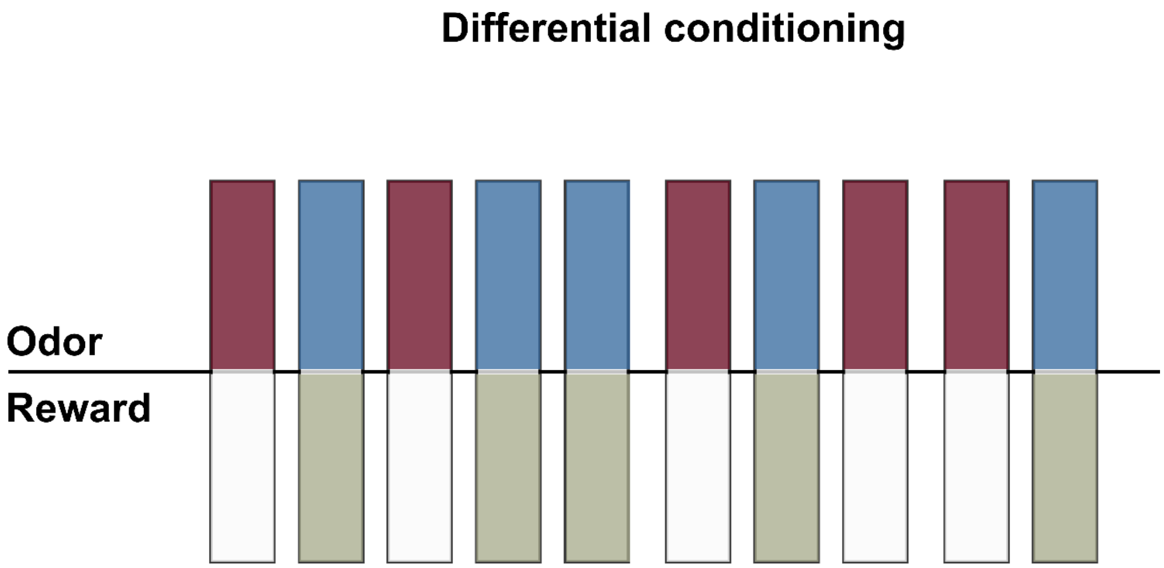

**Supplementary Figure 3. Schematic representation of the generalization conditioning training paradigm, Related to Figure 6a.**

During the training of Exp. 6, honeybees were presented with a total of 14 trials, divided into 8 experimental trials and 6 blank trials. During the blank trials (represented as white bars) a sequence of three puffs of air was provided without any positive reinforcement. In the experimental trials, two odor sequences (ABA or CDC) were presented (first sequence represented as blue bars; second sequence represented as green bars) to the bee in association with a drop of sucrose solution.

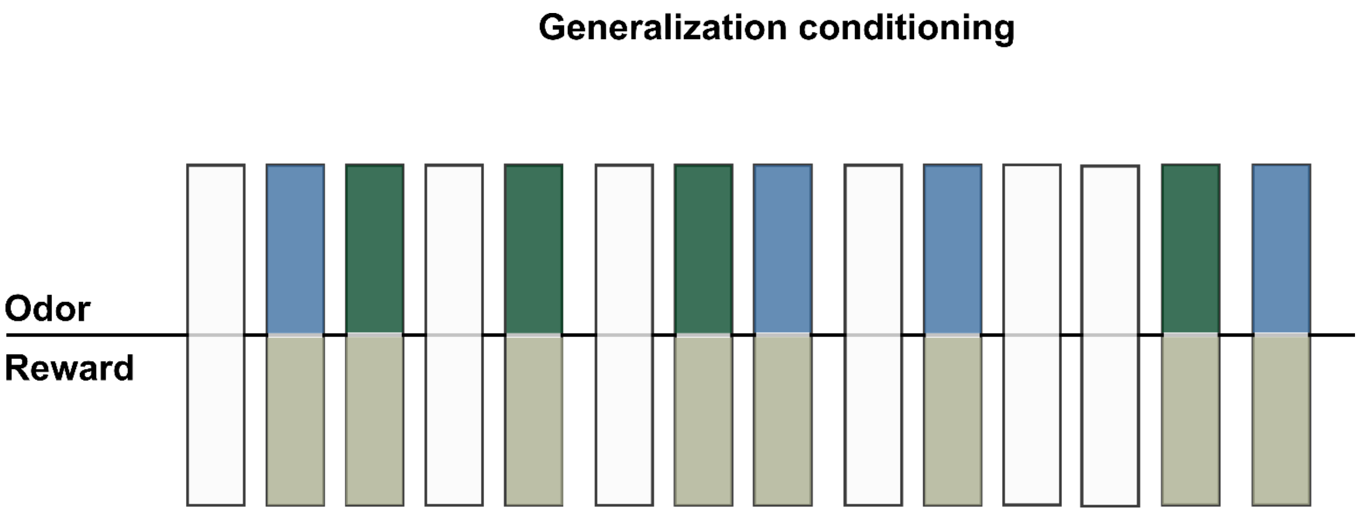

**Supplementary Figure 4. Representation of PER occurrences with respect to the elements of the sequences, Related to Figure 3b and 3c.**

In experiment 3, we encoded the occurrence of the PER response in relation to the presentation of a specific element of the sequence (i.e., first, second, or third element). A similar tendency was observed in both tests (i.e., *memory* test (a) and *recall* test (b)) in which honeybees preferentially responded to the first odor provided, regardless of whether it was the first (A, 3-hexanol) or second (B, acetophenone) element of the previously trained sequence (i.e., ABA).

*(a)*

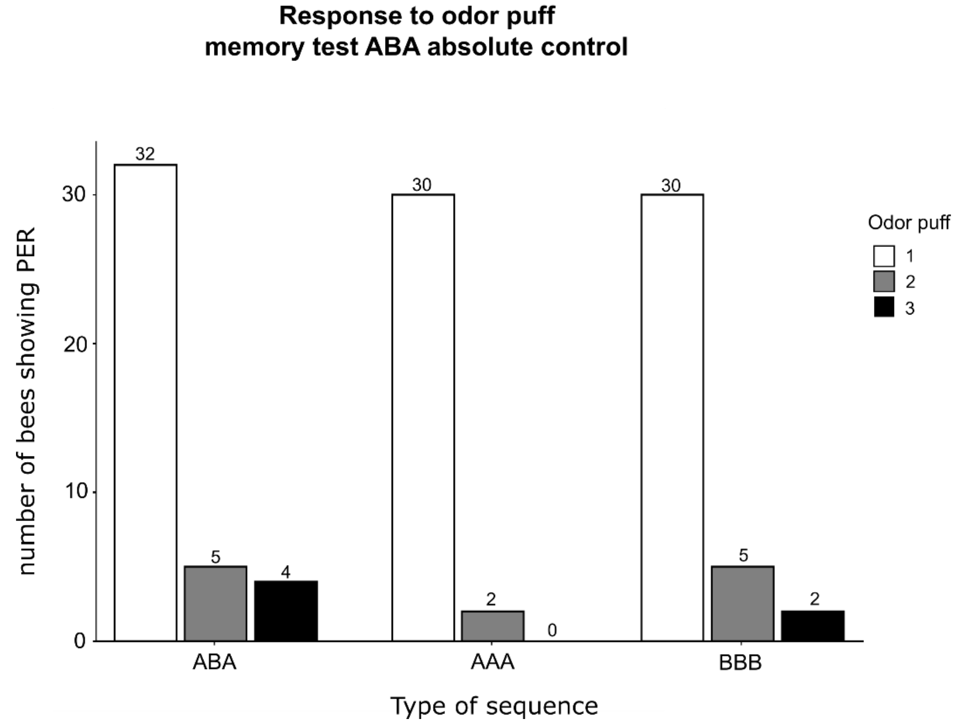


*(b)*

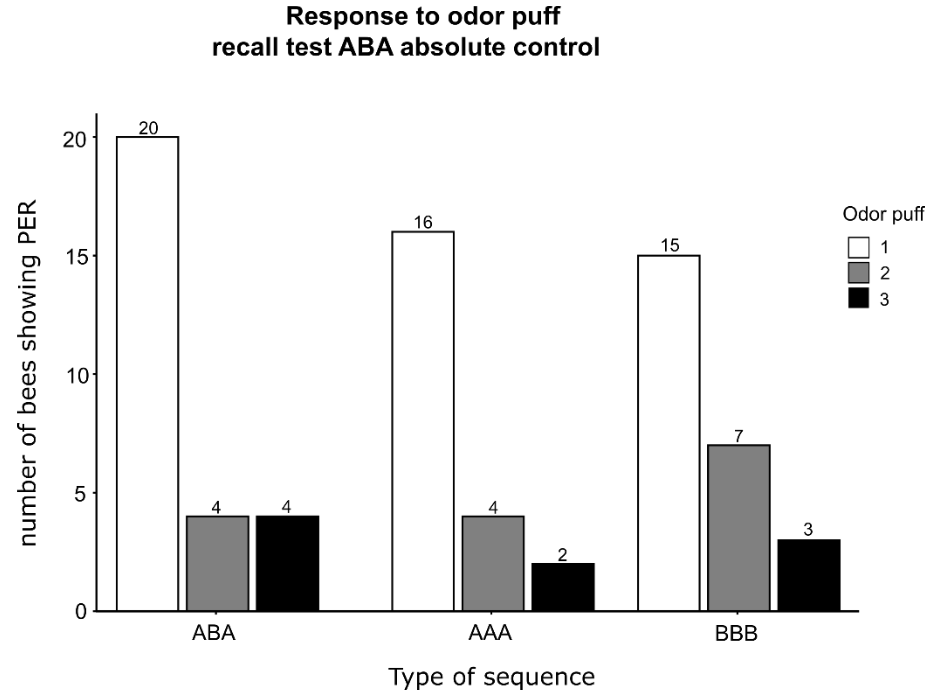

**Supplementary Figure 5. Representation of PER occurrencies respect to the elements of the sequences, Related to Figure 5b and 5c.**

In experiment 5, we encoded the occurrence of the PER response in relation to the presentation of a specific element of the sequence (i.e., first, second, or third element). When either an ABA or AAB sequence was presented (i.e., those were either reinforced or non-reinforced during the training phase), a higher number of bees responded to the odor A (i.e., 3-hexanol) when it occupied the first position in the sequence, compared to when the same odor occupied the second or third position in the sequence. Similarly, when the odor B (i.e., acetophenone) was presented in the second or third position, bees most likely responded to this event. Interestingly, when the response time to novel sequences was scored (i.e., CDC and CCD), the majority of bees responded to the D odor (i.e., citral), irrespective of its position in the sequence. These results were observed both at *memory* test (a) and *recall* test (b).

***(a)***

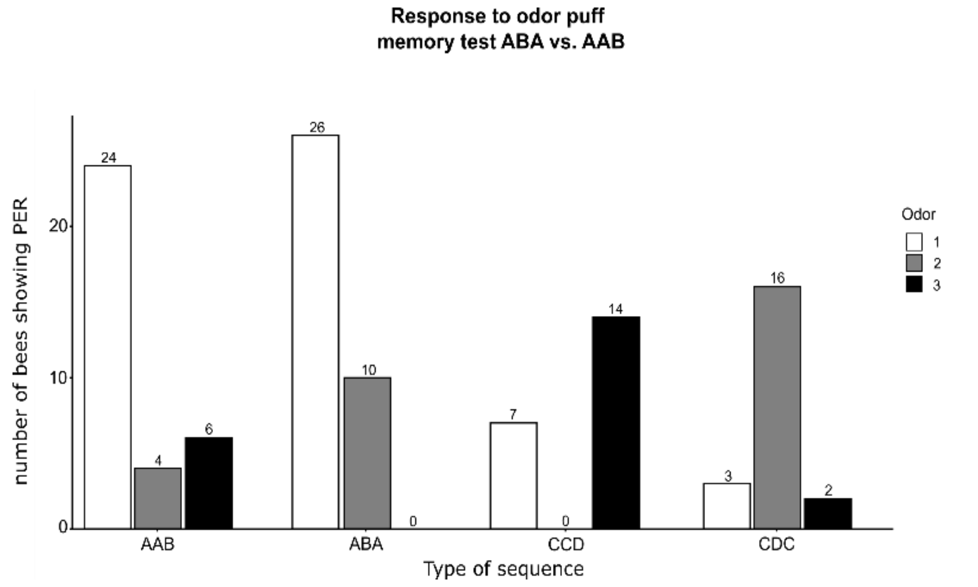


***(b)***

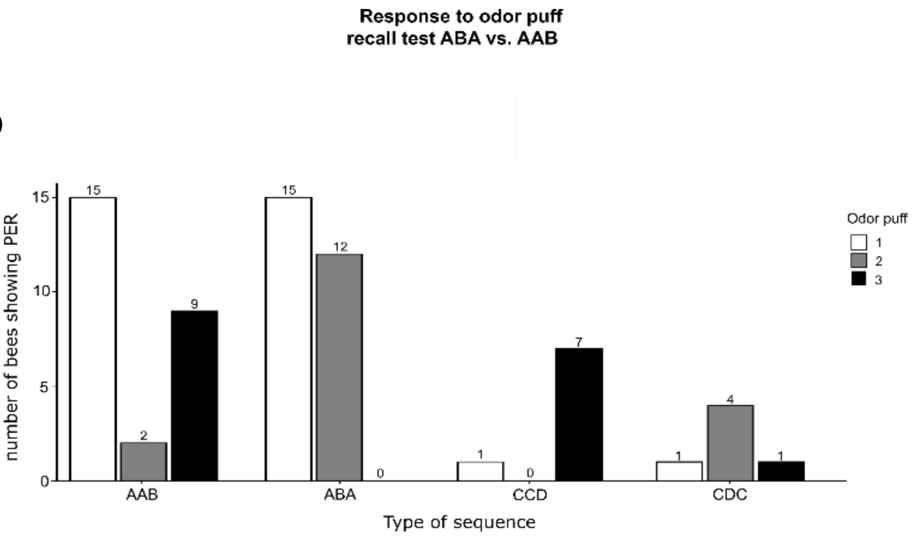
